# Metabolic traits predict the effects of warming on phytoplankton competition

**DOI:** 10.1101/227868

**Authors:** Elvire Bestion, Bernardo Garcia-Carreras, Charlotte-Elisa Schaum, Samraat Pawar, Gabriel Yvon-Durocher

## Abstract

Understanding how changes in temperature affect interspecific competition is critical for predicting changes in ecological communities as the climate warms. Here we develop a simple theoretical model that links interspecific differences in metabolic traits, which capture the temperature-dependence of resource acquisition, to the outcome of pairwise competition in phytoplankton. We parameterised our model with metabolic traits derived from six species of freshwater phytoplankton and tested its ability to predict the outcome of competition in all pairwise combinations of the species in a factorial experiment, manipulating temperature and nutrient availability. The model correctly predicted the outcome of competition in 71% of the pairwise experiments. These results demonstrate that metabolic traits play a key role in determining how changes in temperature influence interspecific competition and lay the foundation for developing theory to predict the effects of warming in complex, multi-species communities.

## INTRODUCTION

Climate change is predicted to be a major cause of species extinctions over the next century (Field *et al.* 2014), and a considerable threat to biodiversity (Bellard *et al.* 2012). Susceptibility to climate change will depend on species’ environmental tolerances (Pacifici *et al.* 2015), with those occupying narrower thermal niches expected to be more vulnerable to climate warming (Magozzi & Calosi 2015). Recent studies have highlighted that changes in species interactions may also play an important role in mediating the impacts of climate change on populations (Dunn *et al.* 2009; Gilman *et al.* 2010; Bellard *et al.* 2012; Cahill *et al.* 2013; Field *et al.* 2014). Indeed, the key drivers of global change (warming, CO_2_ and changes in nutrient availability) are known to affect various types of species interactions, including competition (Tylianakis *et al.* 2008). Understanding how increases in temperatures affect species interactions is therefore crucial to predicting the ecological consequences of future climate change (Dunn *et al.* 2009; Kordas *et al.* 2011; Bellard *et al.* 2012; Dell *et al.* 2014; Reuman *et al.* 2014; Bestion & Cote 2017).

Metabolism shapes numerous life-history traits that determine fitness, including population growth rate, abundance, mortality and interspecific interactions (Brown *et al.* 2004; Savage *et al.* 2004; Dell *et al.* 2011). Species vary widely in the way in which their metabolism and associated ecological rates respond to temperature (Kingsolver 2009; Dell *et al.* 2011). These interspecific differences in thermal performance curves (TPCs) can reflect differences in the magnitude (the elevation of the TPC), sensitivity (relative rate of increase in performance with temperature), and/or thermal optima (the temperature at which the performance is maximised) (Kordas *et al.* 2011; Dell *et al.* 2014; Pawar *et al.* 2015), and can greatly impact species interactions (Reuman *et al.* 2014; Dell *et al.* 2014). Recent theory suggests that differences in metabolic traits between consumers and resources can play a key role in determining the effects of temperature on trophic interactions (Dell *et al.* 2014; Gilbert *et al.* 2014; Pawar *et al.* 2015; Cohen *et al.* 2017). Despite major advances in ecological theory linking the effects of temperature to metabolism and species interactions (O’Connor *et al.* 2011; Dell *et al.* 2014; Gilbert *et al.* 2014; Amarasekare 2015; Uszko *et al.* 2017), there have been very few empirical tests, and to our knowledge, no large-scale experimental study has confronted recent theoretical developments to assess whether differences in metabolic traits between species can predict how interspecific competition responds to warming.

In aquatic ecosystems, temperature and nutrients are the main drivers of phytoplankton productivity (Litchman *et al.* 2010). Phytoplankton exhibit substantial interspecific variation in their responses to temperature and nutrient availability (Eppley & Thomas 1969; Tilman 1981; Aksnes & Egge 1991; Boyd *et al.* 2013; Thomas *et al.* 2016, 2017). These interspecific variations in metabolic and nutrient acquisition traits are widely recognised as being important drivers of competition (Tilman 1981), community assembly (Bulgakov & Levich 1999; Grover & Chrzanowski 2006; Litchman *et al.* 2010; Edwards 2016) and ultimately the productivity of phytoplankton communities (Behrenfeld *et al.* 2005). However, we currently lack experimental tests of theory that can predict the dynamics of competition from differences in metabolic traits between species, which are essential components of models that forecast how the structure and functioning of phytoplankton communities respond to climate change (Follows *et al.* 2007).

Here we address this fundamental knowledge gap by deriving a mathematical model to predict how changes in nutrients and temperature affect the outcome of interspecific competition from differences between species in the metabolic traits that characterize the TPCs of maximum growth rate and performance under nutrient limitation in phytoplankton. We parameterise our model with metabolic traits derived from six freshwater phytoplankton species and test the model’s ability to predict the outcome of competition in all possible pairwise combinations of the six species in a factorial experiment, manipulating both temperature and nutrient availability.

## Theory

We develop a model to predict how interspecific differences in metabolic traits affect the competitive advantage of pairs of competing phytoplankton when both species are rare and colonizing (co-invading) a virgin environment (or patch) (see Section S1 in supporting information (SI) for full model development). This differs from traditional resource competition (Tilman 1981) and adaptive dynamics theory (Dieckman & Law 1996; Diekmann 2003), in that these frameworks assume one competitor (the resident) is at population dynamics equilibrium while the other is introduced into the system at a low density. Here we characterise scenarios where both species are rare and quantify the impact of changes in temperature and resource availability on species’ relative competitive advantage. Because the two populations are initially rare, cells grow exponentially with a constant growth rate and negligible change in nutrient concentration over time. Therefore, before nutrient concentration has been appreciably depleted, population growth rate of the i^th^ species (*i* = *a* or *b*) can be expressed as

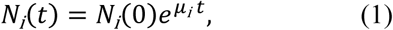

where *N* is the phytoplankton cell density (cells·mL^-1^), *μ* the realised population growth rate (d^-1^), and *t* the time (days). We model growth rate *μ_i_* of the *i*^th^ species using the Monod equation (Monod 1949),

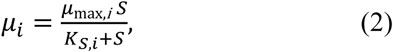

where *μ*_max_ is the maximum growth rate in nutrient-saturated conditions (d^-1^), *K_S_* the halfsaturation constant (pmol-L^-1^), corresponds to the concentration of limiting nutrients at which the growth rate is 50% of *μ*_max_ and measures performance at low nutrient concentrations. *S* is the nutrient (phosphate) concentration (pmol-L^-1^). Maximum growth rate *μ*_max_ is tightly coupled to the rate of net photosynthesis (Geider *et al.* 1998) and consequently, its temperature dependence should follow a left-skewed unimodal function of temperature. Within the ‘operational temperature range’ (OTR, the temperature range typically encountered by the population, see Fig. 1) *μ*_max_ is expected to increase exponentially with temperature (Martin & Huey 2008; Angilletta 2009; Dell *et al.* 2011; Pawar *et al.* 2016). While the temperature-dependence of *K_S_* is less well known (e.g. Carter & Lathwell 1967; Ahlgren 1987; Aksnes & Egge 1991; Sterner & Grover 1998), we assume the same form of temperature-dependence as *μ*_max_ (see Supplementary Section S1 for a discussion of this assumption). We therefore model *μ*_max_and *K_S_* using the Boltzmann-Arrhenius equation (Aksnes & Egge 1991; Reuman *et al.* 2014),

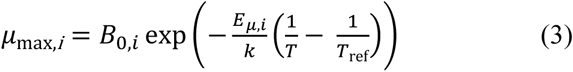

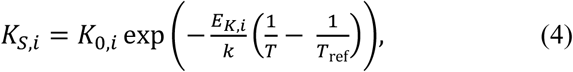

where *B*_0,*i*_ and *K*_0*i*_ are the values of *μ*_max,*i*_ and *K_S,i_* at a reference temperature *T_ref_* (Kelvins) and include the scaling of *μ*_max_ and *K_S_* with cell size (SI section S1), *E_μ,i_* and *E_K,i_* are the activation energies (eV) that phenomenologically quantify the relative rate of change in *μ*_max_ and *K_S_* with temperature, *k* is the Boltzmann constant (eV·Kelvin^-1^), and *T* is the temperature (Kelvins).

**Figure 1.**
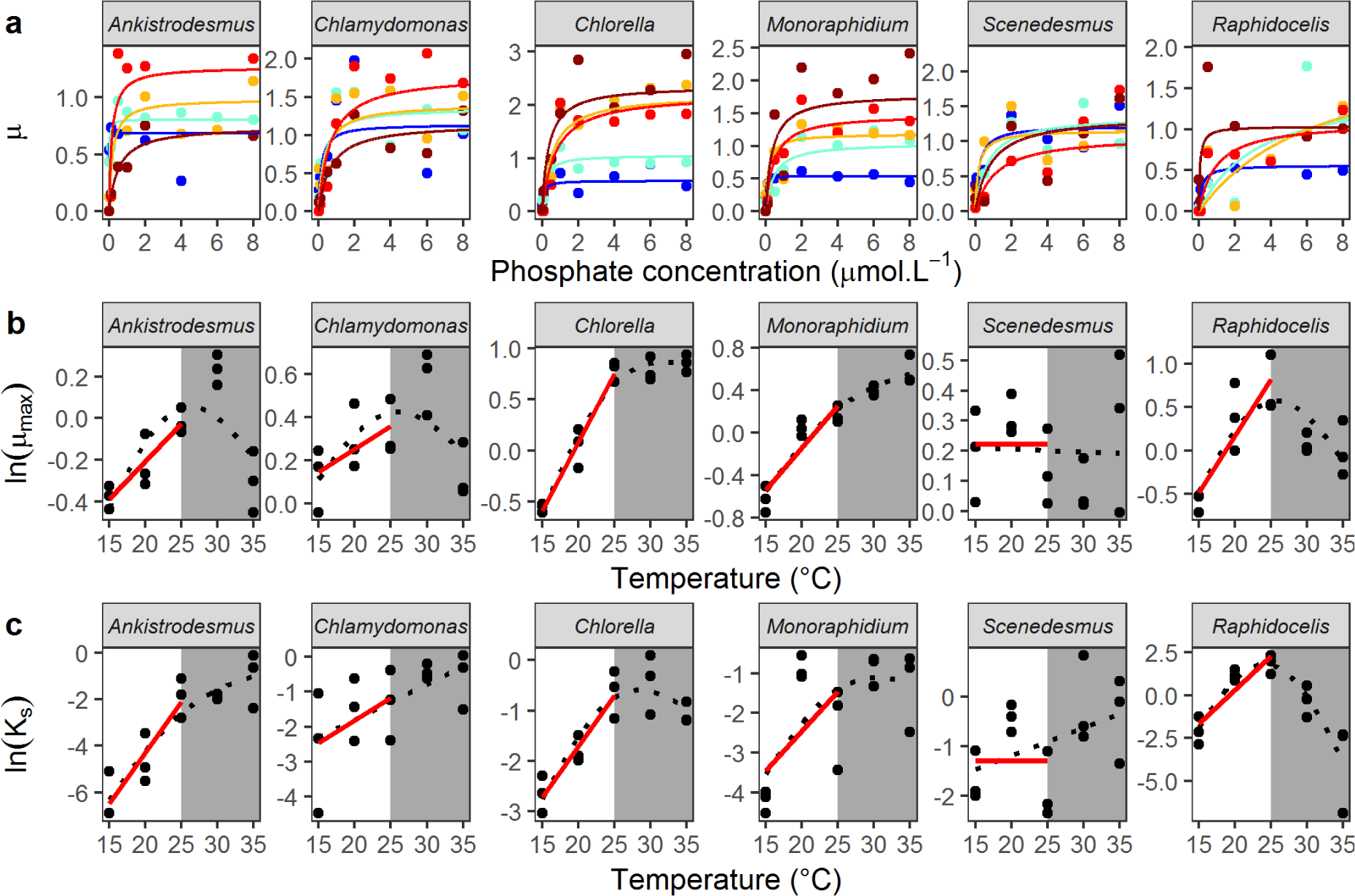
Interspecific variation in metabolic traits. (a) Monod curves for each species, with growth rate *μ* as a function of phosphate concentration (μmol·L^-1^) from 15°C (blue) to 35°C (dark red). Points represent the mean of the 3 replicates, and the Monod curve is drawn from the mean parameters across the 3 replicates. Note that the phosphate concentration levels in the experiment range from 0.01 to 50 μmol·L^-1^ but the x-axis was cut at 8 μmol·L^-1^ for clarity. (b) Maximum growth rate *μ*_max_ and (c) the half-saturation constant *K_S_*, as functions of temperature. Red lines represent the fit of the Boltzmann-Arrhenius within the operational temperature range (15 to 25°C, white area). Black dotted lines represent the fit of the GAM over the whole temperature range. See Tables S4A-D for more details about the temperature dependence of *μ*_max_and *K_S_*.

The parameters of equations (3) and (4) (*B*_0,*i*_,*K*_0*i*_,*E_μ,i_*,*E_K,i_*) are metabolic traits that characterise how resource acquisition and growth respond to temperature.

Assuming *N_a_*(0) = *N_b_*(0) (starting densities are equal in experiments), we can define the competitive advantage (*R*) of species *a* relative to species *b* by taking the log ratio of their abundances at time *t*:

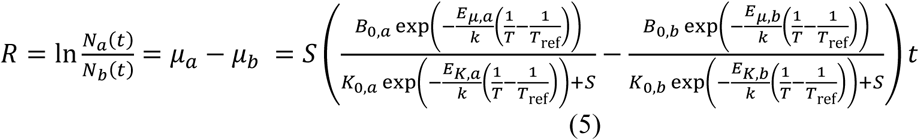

(see SI Section S1). Thus, the value of *R* depends on differences in the competing species’ metabolic traits, that is, on the respective parameters that define the temperature dependence of *μ*_max_ and *K_S_* (*B*_0,*i*_, *E_μ,i_*, *K*_0*i*_, *E_K,i_*) between the two species. When there are no differences (the equivalent parameters are the same in both species), *R* = 0 and both species are expected to be equally abundant at any time point *t*. When there are mismatches, *R* ≠ 0, the sign of *R* indicates which species has a competitive advantage: for *R* > 0, species *a* is expected to outnumber species *b* at time t, while the opposite is true for *R* < 0.

We can assess the relative importance of the metabolic traits characterising nutrient limited and resource saturated growth for predicting competitive advantage by comparing the full model for *R* (equation 5) to a simplified version that assumes nutrient saturation:

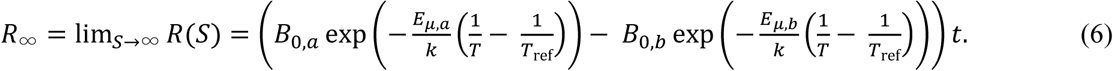

In this case, species *a* will grow faster than species *b* if *R*_∞_> 0, and therefore if

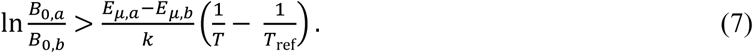

Here, the trade-off between normalisation constants (*B*_0,*a*_, *B*_0,*b*_) and activation energies (*E_μ,a_*, *E_μ,b_*) is explicit. At *T* = *T*_ref_, the winner is determined by the ratio of the normalisation constants (the right hand side of the inequality becomes zero). Species *a* will win when *B*_0,*a*_>*B*_0,*b*_. However, as *T* increases or decreases from *T*_ref_, the relative importance of the activation energies increases, and at sufficiently large |*T* - *T*_ref_|, the winner of the competition is entirely determined by the differences in *E_μ_*. when *T* ≫ *T*_ref_, the winner is the species with the higher *E_μ_* while when *T* ≪ *T*_ref_, the species with the lower *E_μ_* wins (e.g., SI Fig. S1A). For narrower temperature ranges, such as those discussed in this study, the winner is determined by differences in both normalisation constants and activation energies. This tradeoff between the normalisation constants and the activation energies in shaping how the competitive advantage changes with warming is similar (but temperature-specific) to the trade-off functions central to adaptive dynamics.

The sign of *R* and *R*_∞_ can change with temperature — a “reversal” in the competitive advantage indicates that one species can outcompete the other only within a specific temperature range (e.g., Fig. 3; SI Fig. S1B and Section S1). Thus our model makes the following key predictions: (i) differences in individual species’ metabolic traits can predict competitive advantage between pairs of species at a given temperature; (ii) *R*_∞_ will approximate *R* in predictive power at higher nutrient concentrations, but *R* will better predict competitive advantage at lower nutrient concentrations; and (iii) the competitive advantage will reverse with warming if the species with lower performance at low temperature (*B*_0_) has a sufficiently higher thermal sensitivity (*E_μ_*).

**Figure 3.**
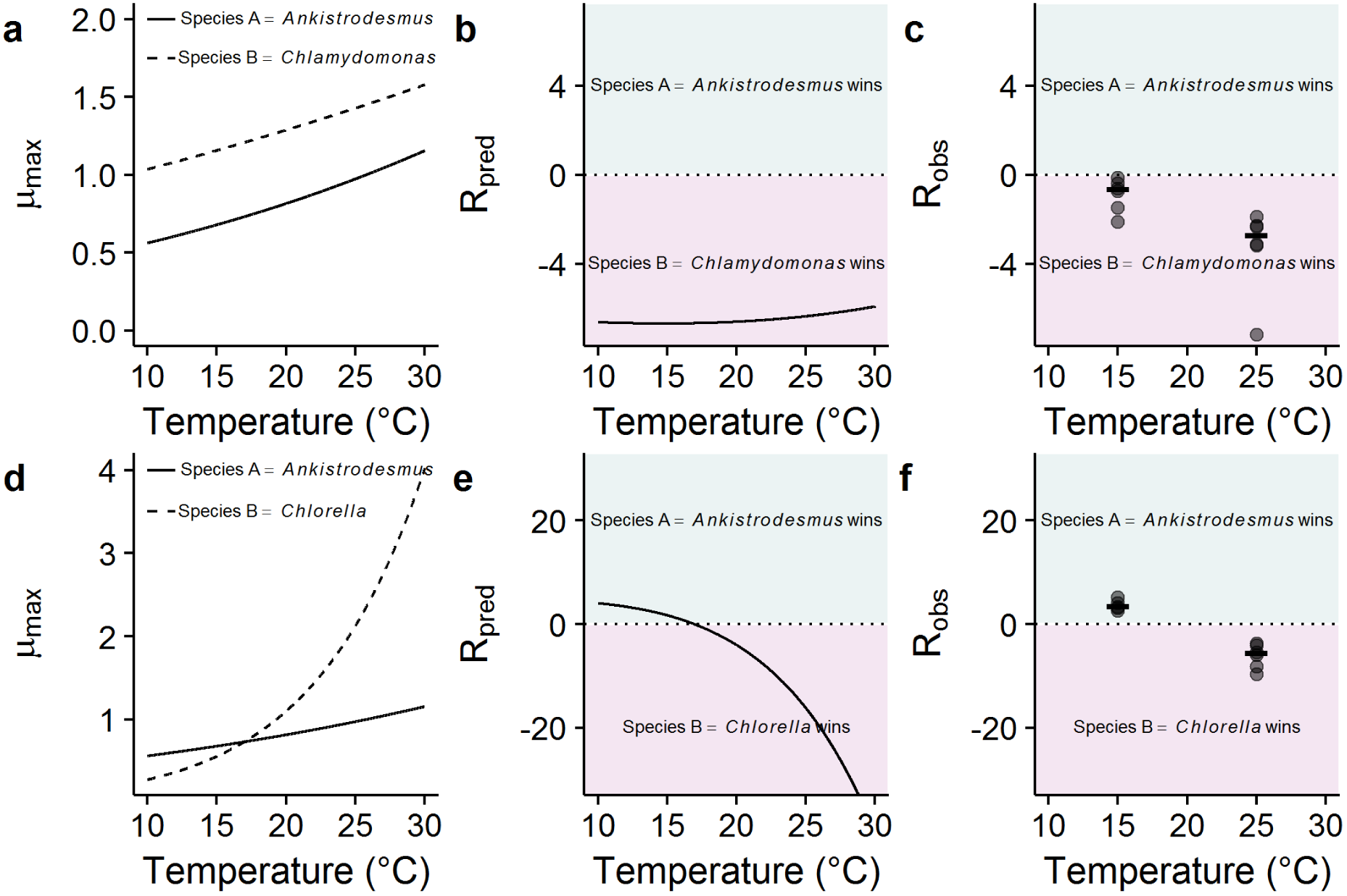
Predicting reversals in competitive advantage from mismatches in metabolic traits. (a-c) competition between *Ankistrodesmus* and *Chlamydomonas,* (d-f) competition between *Ankistrodesmus* and *Chlorella*. (a, d) represent the temperature dependence of *μ*_max_ derived from the Boltzmann-Arrhenius models. In (a), *μ*_max_is always higher for *Chlamydomonas*, while in (d), *Ankistrodesmus* has a higher *μ*_max_ at low temperatures, but a lower *μ*_max_ at high temperatures. This translates into different shapes of predicted *R*_∞_ with temperature, with a reversal of competitive advantage with temperature in the *Ankistrodesmus-Chlorella* competition (e) while there is no reversal in the *Ankistrodesmus-Chlamydomonas* competition (b). These theoretical predictions are in line with the experimental observations (c, f; N = 6 replicates per temperature plus medians as segments).

## METHODS

### Study design

We used an experimental approach to test the model’s ability to predict competition in six phytoplankton species (SI Fig. S2A). We first determined the temperature dependence of *μ*_max_ and *K_S_* for each species independently, which were used to parameterise the model, allowing us to generate predictions on the competitive advantage for each species pair as a function of temperature and nutrient concentration. We then competed the six species in all pairwise combinations at two temperatures and three nutrient concentrations to test the ability of the model to predict the outcome of interspecific competition.

### Species and culture conditions

The experiment was conducted with six species of naturally co-occurring freshwater green algae, *Ankistrodesmus nannoselene, Chlamydomonas moewusii, Chlorella sorokiniana, Monoraphidium minutum, Scenedesmus obliquus* and *Raphidocelis subcapitata* (Fritschie *et al.* 2014). We chose these 6 species because they have similar cell sizes and can be cultured on the same media (standard COMBO culture medium without animal trace elements (Kilham *et al.* 1998)). By choosing similar cell sizes, we aimed to minimize the effect of size on differences in metabolic traits (SI section S1). Strains of each species were ordered in October 2015 from the CCAP (SI Table S2A), and grown on COMBO medium in semi-continuous culture at 15°C on a 12:12 light-dark cycle with a light intensity of 90 μmol·m^-2^.s^-1^, transferring them weekly to keep them in exponential phase of growth until the start of each experiment.

### Metabolic traits

In February 2016, we measured growth rates of each species across gradients in temperature and phosphate concentration. Each species was grown in a factorial experiment at 5 temperatures and 13 phosphate concentrations, with 3 replicates per combination, yielding 1170 cultures (SI Fig. S2A). We created 13 solutions of COMBO medium with different phosphate concentrations ranging from 0.01 to 50 μmol PO_4_^3+^ L^-1^ (SI Table S2B), a range relevant to phosphate concentrations commonly found in lakes (Downing *et al.* 2001). Small tissue culture flasks (Nunclon) filled with 40 mL of each solution were inoculated with each species in monoculture at very low density (100 cells·mL^-1^) ensuring that the increase in phosphate concentration due to the inoculum volume (10 μL) was minimal (0.01 μmol·L^-1^). Cells were then grown at 15, 20, 25, 30, and 35°C, and 90 μmol·m^-2^·s^-1^ on a 12:12 light-dark cycle. Samples were shaken and their position rotated within the incubators daily during the month-long experiment. Every two days, 200 μL was taken and 10 μL of 1% sorbitol solution was added as a cryoprotectant. After one hour of incubation in the dark, samples were frozen at -80°C until further analysis. Cell density was determined by flow cytometry (BD Accuri C6) on fast flux settings (66 μL·min^-1^), counting 10 μL per sample. During the experiment, some samples failed to grow properly and were therefore removed from the subsequent analyses.

### Competition experiments

To investigate the joint effects of temperature and phosphate availability on competition, we competed species in all pairwise combinations (15 pairs) at two temperatures (15 and 25°C; low temperature and a temperature close to the optimum for most species, Fig. 1) and three phosphate concentrations (one saturating [30 μmol·L^-1^] and two limiting [1 μmol·L^-1^ and 0.1 μmol·L^-1^] concentrations, chosen from the Monod curves, Fig. 1), replicated 6 times (SI Fig. S2A), yielding 540 microcosms. We also grew the 6 species in monoculture at the two temperatures and three nutrient levels to train and test an algorithm for discriminating the different species in the competition trials (see SI Section S3 for more details). We used 24 well plates filled with 2 mL of media, inoculated them with 100 cells·mL^-1^ of each species, and incubated them in the same way as described above. After 5, 14 and 23 days, a 200 μL sample was taken and cell density was determined by flow cytometry.

### Data analyses

All statistical analyses were undertaken using R v3.3.2 (R Core Team 2014).

#### Metabolic traits

To characterise the effects of phosphorous availability and temperature on growth we estimated specific growth from the time-series of cell densities. Population dynamics were fitted using non-linear least squares regression to the Buchanan three-phase linear growth model (Buchanan *et al.* 1997):

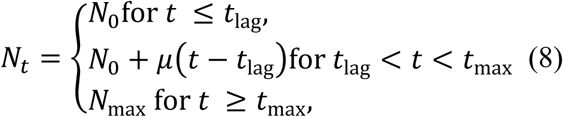

where *t*_lag_ is the duration of the lag phase (days), *t*_max_ the time when the maximum population density is reached (days), *N*_0_ the log_10_ of the initial population density (log_10_(cells·mL^-1^)), *N*_max_ the log_10_ of the maximum population density supported by the environment (log_10_(cells·mL^-1^)), and *μ* the specific growth rate (day^-1^). Fits to the Buchanan model were determined using the ‘nlsLM’ function in the ‘minpack.lm’ package (Elzhov *et al.* 2010), which uses the Levenberg-Marquardt optimisation algorithm. Parameter estimation was achieved by running 1000 different random combinations of starting parameters picked from uniform distributions and returning the parameter set with the lowest AICc score (Padfield *et al.* 2016).

The Monod equation (equation 2, Monod 1949) was fitted to the estimates of *μ* for each species at each temperature and for each of the three replicates using the ‘nlsLM’ function as above.

We used two approaches to describe the thermal variation in *μ*_max_ and *K_S_*, the Boltzmann-Arrhenius model and generalized additive models (GAMs). First, we fitted the Boltzmann-Arrhenius model on ln scales to *μ*_max_ and *K_S_* within the ‘operational temperature range’, between 15 and 25°C, using a reference temperature *T*_ref_ = 15°C (equations 3 and 4) with the ‘nlsLM’ function as above. This analysis produced normalisation constants and activation energies for both *μ*_max_ and *K_S_* per species, which we then used to parameterize equations 5 and 6 in the theory. Second, for each species, we fitted a GAM to ln *μ*_max_ and ln *K_S_* across the full temperature range over which the TPCs are typically unimodal using a basis dimension of 3 and the “ts” type of basis-penalty smoother with the ‘mgcv’ package.

#### Competition

The flow cytometer returned side scatter (SSC), forward scatter (FSC), green (FL1), orange (FL2), red (FL3), and blue (FL4) fluorescence values that can be used to define a species’ morphology and pigment composition. We used these quantities to quantify cell identity and thus estimate the relative abundances of each species in pairwise competition experiments. We separated the data set into three, one for training the discrimination algorithm, one for the testing its efficiency at separating species pairs, and one for the actual competition trials. The training dataset was used to establish pairwise discrimination functions between pairs of species, using three different procedures: a linear discriminant analysis (LDA), a random forest analysis and a recursive partitioning and regression tree analysis (SI Section S3). These different discriminant functions were then applied to the testing dataset to determine the accuracy of the various discrimination algorithms in differentiating between pairs of species by creating *in silico* competition experiments (SI Section S3). The linear discriminant analysis predicted the correct cell identity of each species in the *in silico* pairwise experiments with 78% accuracy and was chosen for application to the competition dataset (SI Fig. S3A and Table S3A). Results were robust to the statistical method used to discriminate between species (SI Section S6).

After determining species identity for each competition trial, we computed cell density and calculated the competitive advantage, R, of species *a* relative to species *b* by taking the ln ratio of their densities (cells·mL^-1^) at time t, and adding one to account for instances when one species had become locally extinct. We also computed a binary competitive advantage where species *a* (respectively species *b*) was competitively dominant for *R* > 0 (respectively *R* < 0).

## RESULTS

### Metabolic traits

The responses of growth rate to phosphate concentration were well fit by the Monod equation (Fig. 1a). The half-saturation constant, *K_S_*, and the maximum growth rate, *μ*_max_, varied with temperature, and the temperature response of these traits differed between species (SI Tables S4A-C). Maximum growth rate exhibited unimodal temperature dependence in *Ankistrodesmus, Chlamydomonas,* and *Raphidocelis* (Fig. 1b, SI Table S4B). In *Chlorella* and *Monoraphidium, μ*_max_ increased with temperature but did not reach a peak by 35°C, while *μ*_max_ in *Scenedesmus* exhibited negligible temperature dependence (Fig. 1b, SI Table S4B). *KS* increased with temperature for *Ankistrodesmus*, *Chlamydomonas*, and *Monoraphidium*, while the response was unimodal for *Chlorella* and *Raphidocelis* and there was no discernible trend for *Scenedesmus* (Fig. 1c, SI Table S4C). The magnitude of the relationship between *μ*_max_ and temperature and between *K_S_* and temperature in the operational temperature range differed between species (Fig. 1b,c, SI Table S4A).

### Interspecific competition

The competitive advantage depended on temperature, nutrient conditions and the identity of the species pair (Fig. 2). For instance, for the pair *Ankistrodesmus-Chlorella, Ankistrodesmus* dominated the competition at lower temperatures while *Chlorella* dominated at higher temperatures, except at very low nutrient concentrations. For some species pairs, one species dominated across temperatures and nutrient concentrations – e.g. *Monoraphidium* always outcompeted *Raphidocelis*. Reversal of the competitive advantage across environmental conditions occurred more often when temperatures changed (in 16 out of 49 competitions) than between shifts in nutrient concentrations (11 out of 22, Fig. 2).

**Figure 2.**
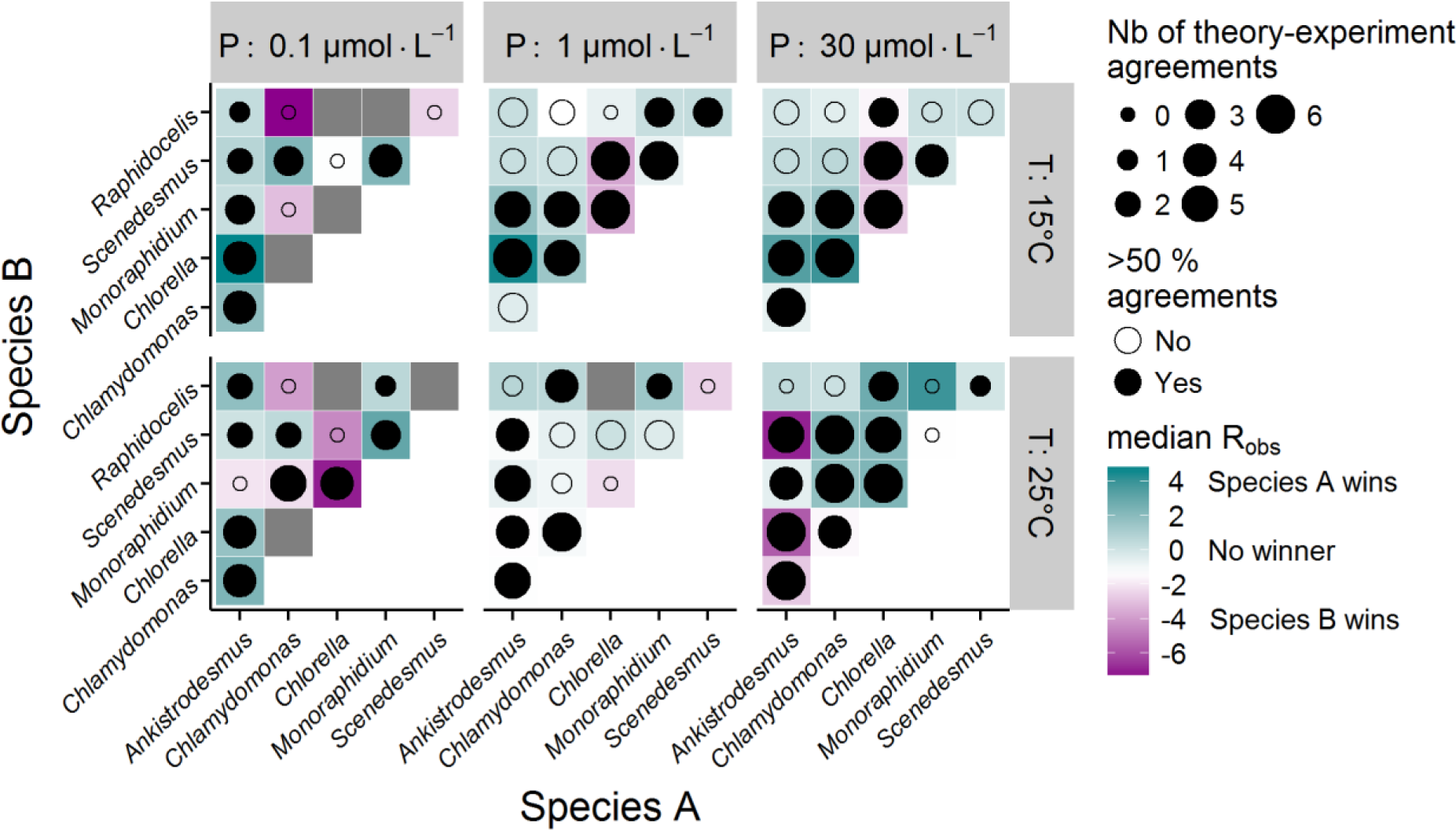
Predicting competitive advantage from metabolic traits. The colour indicates the identity of the competitively dominant species and strength of competitive advantage after 14 days (median *R*_obs_ over 6 replicates; see SI Fig S3B for *R*_obs_ by replicate). The circles show the agreement of the model predictions with the experimental outcomes (size: number of replicates correctly predicted; colour: more than half of the replicates correctly predicted, see Table 1). If the cell density was too low to accurately predict a winner, we dropped the replicate. Thus, the number of replicates per pair, temperature and nutrient conditions is not always 6. 8 competition trials were dropped because all replicates had too low a cell density. These are shown as grey tiles. The total number of replicates is *N* = 369.

The theoretical competitive advantage *R* (equation 5) correctly predicted 71% of the experimental outcomes (Table 1). The predictability of the competitive advantage did not differ between temperatures, but it varied with nutrient concentration and depended on species identity (Table 1). 84% of the interactions involving *Chlorella* were correctly predicted, while those involving *Raphidocelis* were the most difficult to predict (only 48%). Indeed, removing interactions involving *Raphidocelis* increased the overall predictive power of the model to 77%. The model correctly predicted 86% of the observed reversals in competitive advantage across temperatures at the high nutrient conditions, while it was unable to predict reversals at lower nutrient levels (Table 2). These reversals are due to the differences in metabolic traits between species leading to the crossing of growth rate TPCs between two competing species (Fig. 3). Assuming nutrient saturated conditions (*R*_∞_) decreased the predictive power of the model (Table 1). Accounting for interspecific differences in the temperature-dependence of *K_S_* substantially improved predictions at the very low nutrient concentrations.

**Table 1.** Proportion of competitive advantages correctly predicted by theory. Results are shown for the full dataset (including competitions at both temperatures and nutrient concentrations), by temperature, nutrient concentration, and species (where only competitions involving each individual species are considered in turn). The column “*R*_∞_”(equation 6) assumes nutrient saturated conditions, while column “*R*” (equation 5) explicitly captures nutrient limitation. “*N*” indicates the number of competitions in each subset. *P* values indicated in parentheses were obtained by bootstrapping (see SI Section S5). The experimental competition data uses the LDA discrimination method on the results at day 14. Analogous results for the random forest and rpart discrimination methods are shown in SI Tables S6A-B, and for results at day 5 and day 23 are shown in SI Tables S9A-B.

| | $R_{\infty}$ | | $R$ | | $N$ |
| --- | --- | --- | --- | --- | --- |
| *Full dataset* |  |  |  |  |  |
|  | 0.62 | (0.014) | 0.71 | (0.000) | 361 |
| *By temperature* |  |  |  |  |  |
| $T = 15^{\circ}\text{C}$ | 0.66 | (0.069) | 0.71 | (0.012) | 188 |
| $T = 25^{\circ}\text{C}$ | 0.57 | (0.165) | 0.72 | (0.002) | 173 |
| *By nutrient* |  |  |  |  |  |
| $[\text{P}] = 0.1 \mu\text{mol}\cdot\text{L}^{-1}$ | 0.32 | (0.800) | 0.76 | (0.066) | 68 |
| $[\text{P}] = 1 \mu\text{mol}\cdot\text{L}^{-1}$ | 0.64 | (0.994) | 0.66 | (0.000) | 148 |
| $[\text{P}] = 30 \mu\text{mol}\cdot\text{L}^{-1}$ | 0.73 | (0.008) | 0.75 | (0.004) | 145 |
| *By species* |  |  |  |  |  |
| *Ankistrodesmus* | 0.68 | (0.013) | 0.80 | (0.000) | 136 |
| *Chlamydomonas* | 0.61 | (0.048) | 0.70 | (0.006) | 138 |
| *Chlorella* | 0.76 | (0.029) | 0.84 | (0.002) | 119 |
| *Monoraphidium* | 0.60 | (0.067) | 0.72 | (0.009) | 131 |
| *Scenedesmus* | 0.58 | (0.049) | 0.65 | (0.005) | 125 |
| *Raphidocelis* | 0.38 | (0.912) | 0.48 | (0.574) | 73 |

**Table 2.** Number of observed and predicted reversals in competitive advantage between pair of species. Observed reversals are qualified when the median *R* of a pair of species across 6 replicates changes sign with temperature. They are compared to reversals predicted by the model. We counted the number of times the model correctly predicted that a specific pair of species would reverse the sign of their competitive advantage.

|  | *Observed revs.* |  | *Predicted revs. ($R_{\infty}$)* |  | *Predicted revs. ($R$)* |  |
| --- | --- | --- | --- | --- | --- | --- |
|  | *Yes* | *No* | *N* | *Prop.* | *N* | *Prop.* |
| *Full dataset* |  |  |  |  |  |  |
|  | 16 | 23 | 9 | 0.56 | 9 | 0.56 |
| *By nutrient* |  |  |  |  |  |  |
| [P]=0.1 $\mu\text{mole}\cdot\text{L}^{-1}$ | 2 | 8 | 1 | 0.50 | 0 | 0.00 |
| [P]=1 $\mu\text{mole}\cdot\text{L}^{-1}$ | 7 | 7 | 3 | 0.43 | 3 | 0.43 |
| [P]=30 $\mu\text{mole}\cdot\text{L}^{-1}$ | 7 | 8 | 5 | 0.71 | 6 | 0.86 |
| *By species* |  |  |  |  |  |  |
| *Ankistrodesmus* | 7 | 8 | 5 | 0.71 | 4 | 0.57 |
| *Chlamydomonas* | 5 | 9 | 2 | 0.40 | 2 | 0.40 |
| *Chlorella* | 8 | 3 | 6 | 0.75 | 7 | 0.88 |
| *Monoraphidium* | 5 | 8 | 4 | 0.80 | 3 | 0.60 |
| *Scenedesmus* | 5 | 9 | 1 | 0.20 | 1 | 0.20 |
| *Raphidocelis* | 2 | 9 | 0 | 0.00 | 1 | 0.50 |

The observed competitive advantages between species pairs were correlated between day 5 and day 14 (Pearson r = 0.49 [0.35, 0.60]) and between day 14 and day 23 (r = 0.49 [0.39, 0.57]). Furthermore, the performance of the model in predicting the competitive advantage was also consistent between time points, with the model correctly predicting 67% of interactions after 5 days, 71 % after 14 days and 68 % after 23 days (Section S9).

We also tested the model’s ability to quantitatively predict the magnitude of *R*. We found a significant correlation between the predicted and observed *R* (SI Fig. S7A, Table S7A), which became stronger when excluding *Raphidocelis* (SI Tables S7B-C).

## DISCUSSION

Understanding how changes in temperature and nutrients affect competitive interactions among phytoplankton is critical to predicting how environmental change will shape the structure and functioning of aquatic ecosystems. We addressed this challenge by developing, parameterizing and testing a model that predicts competition among phytoplankton from differences in the traits that characterize the TPCs of maximum growth rate and performance under nutrient limitation. Our analyses demonstrate that the relative competitive advantage of six species of freshwater phytoplankton under changing temperatures and nutrients can be predicted with information on just four metabolic traits.

In our experiments, the response of growth rate to phosphorous availability was well fit by the Monod equation. The parameters characterizing this functional response to resource availability were temperature dependent. Over a broad range of temperatures (15 to 35°C) both the maximum growth rate (*μ*_max_) and the half saturation constant (*K_S_*) exhibited nonlinear temperature-dependence, consistent with Senft *et al.* (1981). However, within the operational temperature range (OTR), the temperature dependence of both *μ*_max_ and *K_S_* were well fit by the exponential Boltzmann-Arrhenius equation. This result is interesting *per se* as, compared to *μ*_max_, the temperature-dependence of *K_s_* is poorly understood (see Section S1). Our results support the positive temperature dependence expected by some theoretical studies (Goldman & Carpenter 1974; Aksnes & Egge 1991; Reuman *et al.* 2014). For both *μ*_max_ and *K_S_*, the activation energies and normalisation constants (value of the trait at a reference temperature) differed among the six phytoplankton species.

We used these empirically determined metabolic traits to parameterize our model to predict the effects of changes in temperature and nutrients on the relative competitive advantage of each species in competition with each of the others and tested the outcome against a factorial experiment, manipulating temperature and nutrient availability. Our experiment revealed that species’ relative competitive advantage changed substantially with temperature and nutrients. Comparing the model’s predictions to the experimental results demonstrated that differences in metabolic traits were a good predictor of the relative competitive advantage of a species in pairwise competition, with the full model correctly predicting 71% of the experimental outcomes. Accounting for the effects of temperature on nutrient limited growth kinetics (R) was important for predicting species’ competitive advantages under very low nutrient concentrations (0.1 μmol PO_4_^3+^ L^-1^), but as nutrient concentration increased, knowledge of differences in the temperature dependence of *μ*_max_ were sufficient to predict the effects of warming at intermediate (1 μmol PO_4_^3+^ L^-1^) and high (30 μmol PO_4_^3+^ L^-1^) nutrient concentrations.

For some combinations, one species was dominant at all temperatures and nutrient concentrations. In these cases, the competitively superior species often had a higher normalisation constant for maximum growth rate (i.e., *B*_0_) resulting in faster realized growth rate under all conditions (Fig. 3). There were also frequent reversals of competitive advantage, particularly with changes in temperature. Temperature-driven reversals in competitive advantage were often linked to analogous reversals in the competitive advantage predicted by the model, where the superior competitor in the warm environment typically had a higher activation energy for maximum growth rate (*E_μ_*, Fig. 3). The model predicted 86% of competitive reversals at high nutrient levels. The poor predictability at low nutrient concentrations may simply reflect the fact that temperature-driven competitive reversals were generally rare (n = 2). Indeed, the model’s overall performance under nutrient limited conditions was very good, predicting outcomes in 76% of cases. The lack of temperature-driven reversals in competitive advantage under nutrient limited conditions suggests that normalisation constants for *μ*_max_ and *K_S_* were the main drivers of competition rather than the activation energies, perhaps because, the temperature dependence of growth and resource uptake are heavily constrained at low nutrients (Thomas *et al.* 2017). Overall, these results demonstrate that metabolic traits play a central role in shaping competitive interactions among phytoplankton and highlight that particular combinations of traits consistently predict competitive advantage under warming – i.e., high *B*_0_ and *E_μ_*. Our findings also suggest that a greater understanding of the variation in metabolic traits at local to global scales is urgently needed if we are to predict how the structure and functioning of planktonic ecosystems will be affected by climate change (Litchman & Klausmeier 2008; Litchman *et al.* 2010).

Despite the good agreement between our model and the median experimental outcomes, the results should be interpreted with some caution because the experimental competitive coefficients were often variable among the six replicates in each pairwise interaction (Figure S3B). Such variability might reflect natural intra-population variability in traits, which is not captured by the model that is parameterized by the average trait values for each species. It could also be driven by experimental precision in quantifying the competitive advantage in small volume, high-throughput batch-culture experiments. Future work will be needed to verify these results in smaller scale experiments using high precision chemostat methods. Nevertheless, the competitive advantages were generally highly predictable, particularly when excluding interactions involving *Raphidocelis*, suggesting that the model’s assumptions are nonetheless appropriate for the other five species. The poor predictability of interactions involving *Raphidocelis* warrants further attention. Our ability to discriminate and quantify this species when in competition using the linear discriminant algorithm was poor (Table S3A), and the confidence intervals around the TPCs of *μ*_max_ and *K_S_* were also wide (Fig. 1, SI Tables S4A-C), which might have impaired the performance of the model. Other factors not accounted for in the model, such as direct interspecific interference (e.g., through the production of toxins), might be more important in this species’ interactions. Indeed, total polyculture yields involving *Raphidocelis* were substantially lower than expectations based on the weighted average of the monoculture yields (Table S8A, Loreau & Hector 2001), indicating strongly negative interactions that would be consistent with interspecific interference.

Our experiments and theory explored the short-term dynamics of two species colonising virgin environment when both are locally rare. The model can however also be extended to explore scenarios, where a rare species (or genotype) invades a resident that is at population dynamics equilibrium (see SI Section S1), which are central to resource competition theory (Tilman 1981), modern coexistence theory (Chesson 2000), and “adaptive dynamics” (Dieckman & Law 1996; Diekmann 2003). Tilman *et al.* (1981) proposed that the outcome of competition is determined by the species with the lowest *R\** (in our notation, *S\**), that is, the species with the lowest equilibrium resource requirements. The *R\** could, for this purpose, be derived from our model with the explicit temperature-dependent parameters we use here (*μ*_max_,*K_S_*), leading to predictions for the effects of differences in metabolic traits on invasion under a range of warming and nutrient manipulation scenarios. Tilman *et al.*’s *R\** concept also extends to the adaptive dynamics framework, where the difference in *R\** between a resident and a competing genotype is equivalent to the ‘invasion fitness’ criterion (e.g., see Section 4 in (Diekmann 2003)). As with resource competition theory, for a competing genotype to successfully invade, its *R\** needs to be lower than that of the resident. Differences in the temperature dependence of species’ metabolism (or those of residents and mutants) would therefore be expected to lead to trade-offs in invasion fitness, comparable to those we have observed in the context of temperature-driven reversals in competitive advantage owing to species’ differences in the activation energy and normalisation constant of maximum growth rate and the half saturation constant (see Fig. 3).

A key assumption of our model is that populations are initially rare and cells grow exponentially with a constant growth rate and negligible change in nutrient concentrations over time. This assumption was violated in several of the experimental conditions at day 14, which were the data used to test the theory. The median time to carrying capacity during the single species nutrient-gradient experiments at 15 and 25°C, were 11 and 9 days respectively at 0.1 μmol PO_4_^3+^ L^-1^, 11 and 7 days at 1 μmol PO_4_^3+^ L^-1^, and 15 and 9 at 30 μmol PO_4_^3+^ L^-1^. At high temperatures and low nutrient concentrations many species were no longer in the exponential phase of growth. We assessed the impact that this violation in the model’s assumptions might have on the model-data comparisons by quantifying the correlation between the competitive advantages derived at day 5 (when all species were still in exponential growth under all conditions) with those used to test the model at day 14. Competitive advantages were strongly correlated between day 5 and 14 (as well as between day 14 and 23). These results demonstrate that the competitive advantage at day 14 carries the signature of exponential growth suggesting that because the initial competitive advantage results in an exponentially higher abundance of the competitively superior species (SI equations (15) and (20)), this advantage persists well into the stationary phase of community assembly even if the species are no longer growing exponentially at the point of sampling.

Overall, our study shows that temperature-driven shifts in competitive advantage among phytoplankton can be predicted from basic information on the metabolic traits governing the thermal responses of growth and resource acquisition. These results emphasize the potential for using metabolic traits to predict how directional environmental change (e.g., climatic warming) as well as environmental fluctuations influence the ecological dynamics of phytoplankton communities. Extending our theoretical and empirical work beyond pairwise interactions to complex multi-species communities will require further work in two main areas. First, the theory will need to be extended to understand how differences in metabolic traits play out in the context of indirect interactions in multi-species trophic interaction networks (Wootton 1994; Menge 1995; Montoya *et al.* 2009). Second, a more comprehensive understanding of metabolic trait variation at local and regional scales will be needed to expand the pairwise models to a trait-based meta-community framework for the effects of climate change on community dynamics.

## Acknowledgements

We thank Saskia Johnson and Emily Budd for their help in the preliminary experiments. This work was supported by a NERC standard grant awarded to SP and GYD (NE/M003205/1; NE/M004740/1).

